# Exocytosis of the silicified cell wall of diatoms involves extensive membrane disintegration

**DOI:** 10.1101/2021.09.14.460270

**Authors:** Diede de Haan, Lior Aram, Hadas Peled-Zehavi, Yoseph Addadi, Oz Ben-Joseph, Ron Rotkopf, Nadav Elad, Katya Rechav, Assaf Gal

## Abstract

Diatoms are unicellular algae, characterized by silica cell walls. The silica elements are formed intracellularly in a membrane-bound silica deposition vesicle (SDV), and are exocytosed after completion. How diatoms maintain membrane homeostasis during the exocytosis of these large and rigid silica elements is a long-standing enigma. We studied membrane dynamics during cell wall formation and exocytosis in two model diatom species, using live-cell confocal microscopy, transmission electron microscopy and cryo-electron tomography. Our results show that during the formation of the mineral phase it is in tight association with the SDV membranes, which are forming a precise mold of the delicate geometrical patterns. During exocytosis, the distal SDV membrane and the plasma membrane gradually detach from the mineral and disintegrate in the extracellular space, without any noticeable endocytic retrieval or extracellular repurposing. Within the cell, there is no evidence for the formation of a new plasma membrane, thus the proximal SDV membrane becomes the new barrier between the cell and its environment, and assumes the role of a new plasma membrane. These results provide direct structural observations of diatom silica exocytosis, and point to an extraordinary mechanism in which membrane homeostasis is maintained by discarding, rather than recycling, significant membrane patches.

**Significance Statement:** Exocytosis is a fundamental process for cell metabolism, communication, and growth. During exocytosis, an intracellular vesicle fuses with the plasma membrane to release its contents. In classical exocytosis, where the exocytosed vesicles are much smaller than the cell, membrane homeostasis is maintained by recycling excess membranes back into the cell. However, an extreme case of exocytosis is the extrusion of large and rigid cell wall elements by unicellular marine algae. During this process, the cell needs to deal with a potential doubling of its plasma membrane. This study reports on a unique exocytosis mechanism used by these organisms, in which the cells cope with the geometrical and physical challenges of exocytosis by releasing a significant amount of membranes to the extracellular space.

## Introduction

Diatoms are a diverse group of unicellular algae characterized by silica cell walls with intricate, species-specific shapes and hierarchical pore patterns (1). Despite immense morphological diversity between species, most diatom cell walls have a conserved layout of two similarly shaped silica ‘shells’ that partially overlap, like a petri-dish. Each ‘shell’ in itself consists of a valve and a series of girdle bands. The valves usually define the shape of the cell, are richly ornamented, and contain hierarchical pore patterns. The girdle bands form partially overlapping rings that surround the sidewalls of the cells.

Diatom cell wall formation is under biological control and linked to the cell cycle (2–5). Each daughter cell inherits one half of the parental cell wall and forms a second valve directly after cell division. New girdle bands are formed and appended to the new valve during the growth of the cell (Fig. S1). Silicification is usually an intracellular process, taking place inside a membrane-bound organelle, the silica deposition vesicle (SDV) (6–8). SDVs are elongated, but very thin organelles, positioned directly under the plasma membrane. Inside the SDV, the chemical environment and biological regulation exert tight control over the silicification process, resulting in highly specific and reproducible cell wall architectures (9–13). After intracellular maturation, the silica elements are exocytosed to cover the cell surface (6, 14). Such exocytosis events are exceptional in cell biology since the content of the SDV is an enormous solid structure that needs to be secreted without damaging the integrity of the cell.

In classical exocytosis pathways, the membrane of an intracellular vesicle fuses with the plasma membrane to deliver its content into the extracellular space. Homeostasis of the plasma membrane can be maintained through a transient fusion that prevents the secreting vesicle from integrating with the plasma membrane, or by offsetting the added membrane through compensatory endocytosis (15). However, due to their large size and rigidity, the secretion of diatom cell walls inflict a huge challenge to the organism. The new valve covers as much as half of the total cell surface and thus the surface area of the SDV membrane is similar to the entire plasma membrane, therefore, fusion with the SDV membrane would lead to an almost instantaneous doubling of the plasma membrane area.

While former studies have brought forward several hypotheses for silica cell wall exocytosis in diatoms (14, 16–23; Fig. S2), significant experimental challenges have hindered further progress. On the one hand, traditional sample preparation for ultrastructural studies with electron microscopy causes artefacts like membrane shrinking and structure deformation (24). On the other hand, the advantages of light microscopy in studying the dynamic aspects of cell wall formation in living cells are limited by the low spatial resolution and scarcity of molecular biology tools for diatoms (25, 26). Due to these limitations, in situ observations of the SDV in the cellular context are sparse, and direct evidence for the nature of the silicification and exocytosis process is missing.

In this study, we investigated membrane dynamics during valve formation and exocytosis in two model diatom species, *Stephanopyxis turris* and *Thalassiosira pseudonana*, using live-cell confocal fluorescence microscopy, transmission electron microscopy (TEM) and cryo electron tomography (cryo-ET). The relatively large *S. turris* cells have easily discernible silica structures, allowing detailed observations of individual valves and the dynamic development of their architectural features using fluorescence microscopy (27, 28). In addition, using cryo-ET we obtain high-resolution 3D data of the SDV in *T. pseudonana*, at native-like conditions (29, 30). Our results indicate that valve exocytosis in both species involves disintegration of the distal membranes, accompanied by the repurposing of the proximal SDV membrane into a new plasma membrane.

## Results

We investigated valve exocytosis in diatoms by studying two species, *S. turris* and *T. pseudonana* (Fig. 1). The valves of *S. turris* are capsule-shaped and have a two-layered structure. The proximal layer is thin, perforated by nano-sized pores, and on the distal side overlain by a more elevated layer that forms large polygons. The top of the polygonal layer is flattened, forming a ‘T’-shape in cross-section (Fig. 1 A’’). *S. turris* cells form chains that are linked through tubular linking extensions of silica that extend from the apex of the valves (Fig. 1 A, arrow). *T. pseudonana* cells are shaped as a barrel and are much smaller, the valves are discs with a diameter of about 5 μm (Fig. 1 B). Small pores perforate the silica layer that spans the area between radial ribs, and larger tubular pores, called fultoportulae, decorate the rim of the valve (Fig. 1 B’) (7). Figure 1 E and F show dividing cells during and shortly after valve formation. Newly formed valves are stained by PDMPO, a fluorescent dye that is incorporated into silica during the process of biological mineralization (31). Comparing the cell size to the size of mature valves shortly before exocytosis illustrates the enormous challenge involved in exocytosis of these rigid silica cell walls (Fig. 1 G, H).

**Figure 1.**
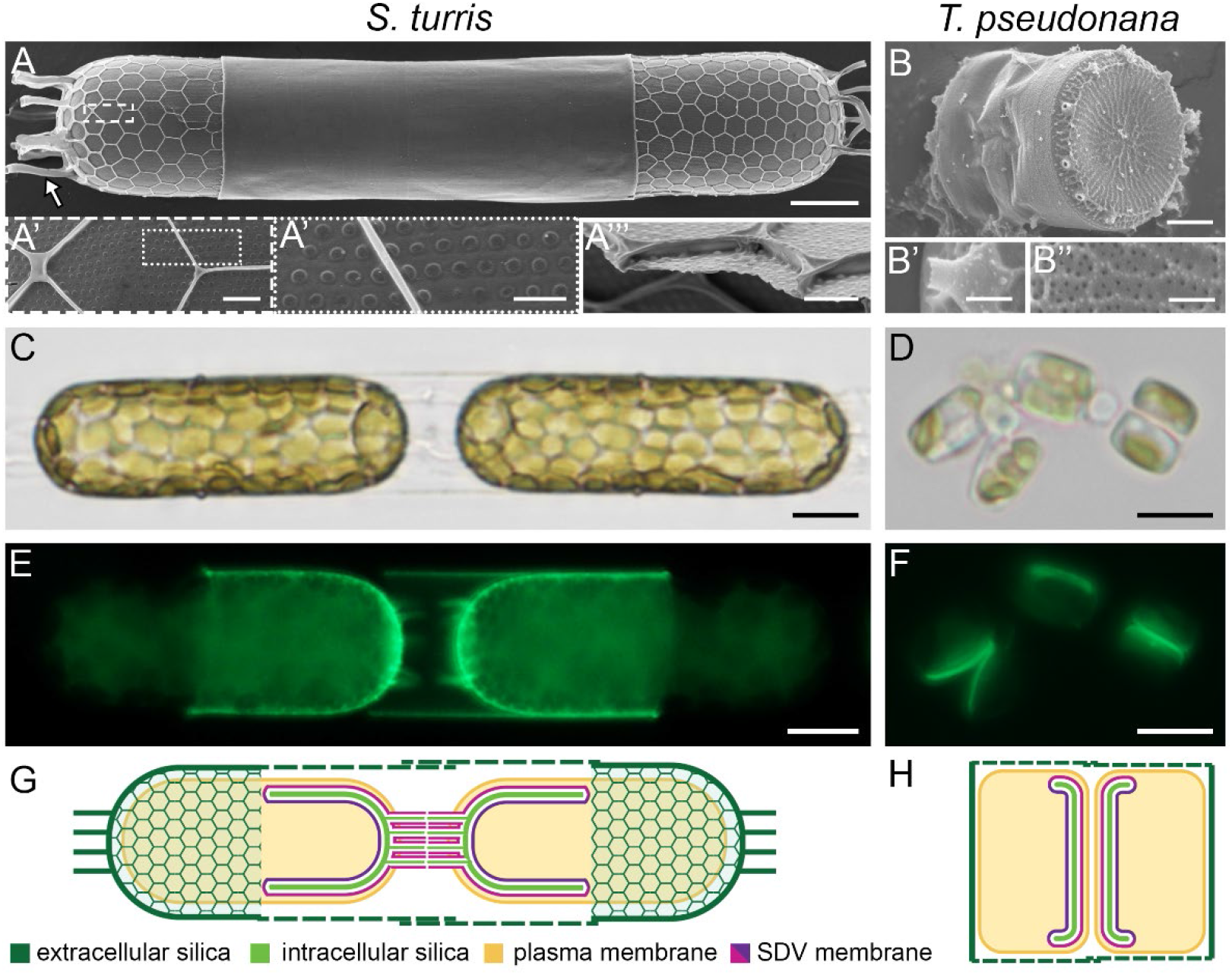
Cell architecture of *S. turris* (A, C, E, G) and *T. pseudonana* (B, D, F, H). (**A-B**) SEM images. Details of the silica valve structures are shown at higher magnifications in the sub-panels. (**C-D**) Bright field light microscope images of two *S. turris* cells in a chain connected through linking extensions and several *T. pseudonana* cells at different stages during the cell cycle. (**E-F**) PDMPO fluorescence image of two different *S. turris* daughter cells during valve formation, and the same *T. pseudonana* cells as in (**D**) with new valves fluorescently stained by PDMPO. (**G-H**) Schematic of *S. turris* and *T. pseudonana* depicting the situation shortly before exocytosis of intracellularly formed valves. Scale bars: 10 μm (**A, C, E**), 5 μm (**D, F**), 1 μm (**A’, A’’’, B**) 500 nm (**A’’**), 100 nm (**B’, B’’**).

We studied membrane dynamics in *S. turris* using time-lapse confocal microscopy on living cells (Fig. 2). Synchronized cultures were labelled with PDMPO to track silica formation (Fig. S3), and with FM4-64 to stain the plasma membrane. FM4-64 is an amphiphilic, membrane-impermeable dye, that enters the cell only via endocytic vesicles (32, 33). During silicification, the PDMPO and FM4-64 signals are almost indistinguishable, demonstrating the close proximity of the plasma membrane to the growing silica structures within the SDV (Fig. 2 A). This is exemplified during the formation of the polygons and the linking extensions, for which the fluorescent membrane signal by itself mirrors the process of silica growth. Nevertheless, the growing tips of linking extensions are exclusively stained by the membrane dye, indicating that SDV elongation precedes silicification (Fig. 2 A, arrow).

**Figure 2.**
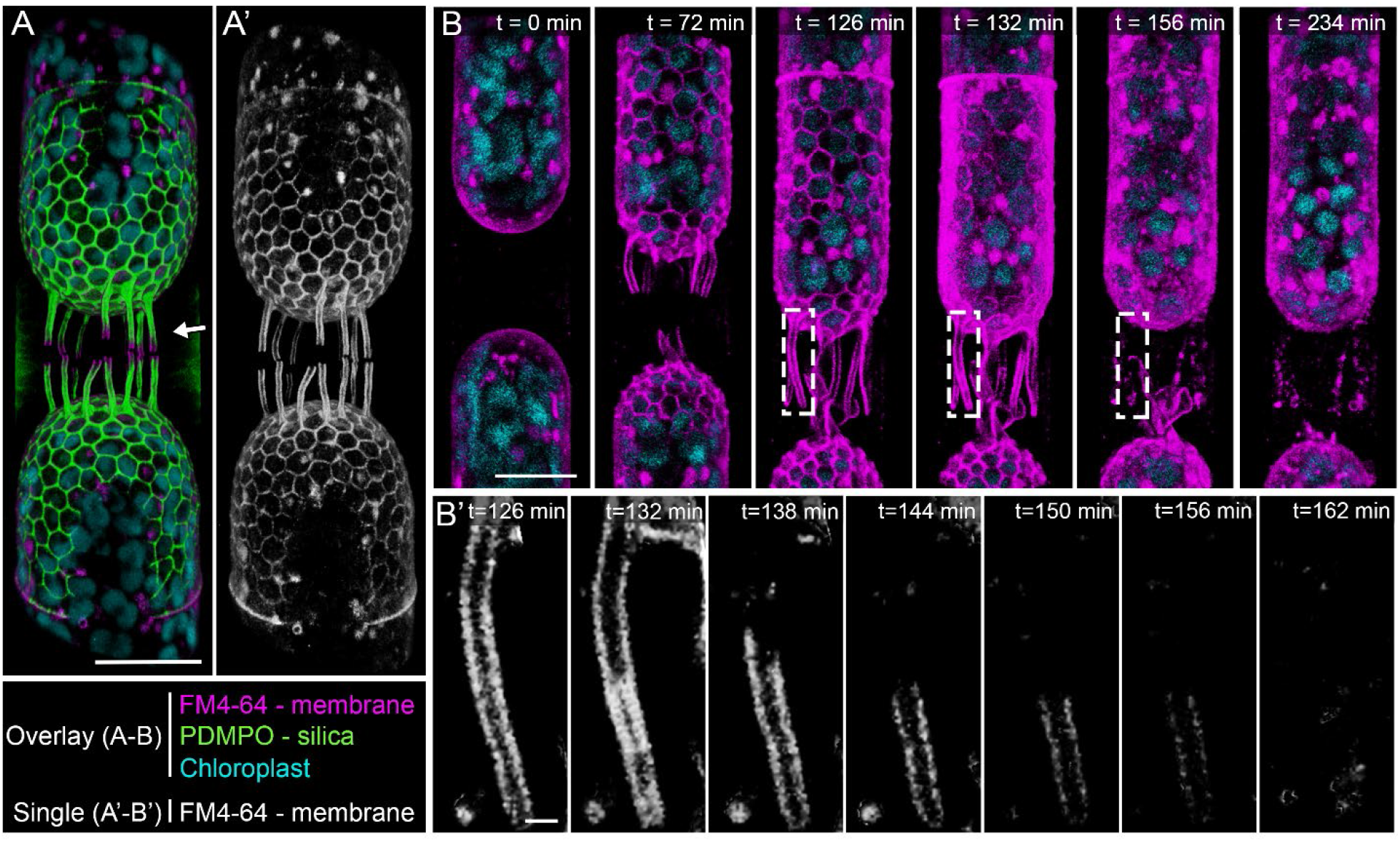
3D reconstructions of time-lapse confocal fluorescence images showing coordinated silica and membrane dynamics during valve formation in *S. turris*. **A, B**) Overlay images of fluorescence channels. **A’, B’**) Single channel FM4-64 fluorescence images. (**A-A’**) Daughter cells during valve formation. Membranes outline the growing polygons and lead the growth of linking extensions (arrow). (**B**) Snapshots from a time-lapse of daughter cells going through valve formation and exocytosis. Elapsed time since start of imaging is indicated in the image. (**B’**) Cropped and magnified view of a single linking extension of the cell in **B** (indicated with boxes). Onset of exocytosis of the valve can be seen from t=132, after which the membrane remnants around the linking process are no longer connected to the cell and gradually disintegrate. Scale bars: 20 μm (**A, A’**), 10 μm (**B**), and 1 μm (**B’**).

We recorded a time-lapse of the membrane dynamics in *S. turris* cells going through the entire process of valve formation and exocytosis (Fig. 2 B, Movie S1). Silicification starts at the cell apex and advances radially. Progress of valve formation is visualized by the fluorescent plasma membrane outlining the growing polygonal silica layer and linking extensions (Fig. 2 B, t=0 to t=126). The onset of exocytosis, namely the loosening of the plasma membrane – SDV – silica complex is evident as the labelled plasma membrane no longer sharply outlines the polygons (Fig. 2 B, t=132). At the same time point, we observe an enhancement of the fluorescent signal (Fig. S4), likely due to fusion between the plasma and SDV membranes that exposes the SDV lumen to the surrounding media, from which free FM4-64 can infiltrate and stain the SDV membrane additionally to the plasma membrane. Nevertheless, the limited resolution of fluorescence microscopy does not allow to spatially resolve the interplay between SDV and plasma membranes before and after exocytosis (Fig. S5).

The linking extensions of *S. turris* present a unique opportunity to investigate the exocytosis process as the old plasma membrane surrounds these long structures before exocytosis, but after exocytosis, the new plasma membrane is only at their base. The FM4-64 signal around the linking extensions is continuous during their growth, but after the extensions reached their full size the fluorescent signal changes to interrupted patches that gradually disappear (Fig. 2 B, t=156 and t=234). In some cases, the membranes surrounding a linking extension are clearly no longer connected to the cell body (Fig. 2 B’). This labelling pattern, in which patches of stained membranes are disconnected from the cell, is very different from the expectation for a classical exocytosis process where stained membranes should be withdrawn and recycled inside the cell. It is important to note that FM4-64 is only transiently attached to the membranes and is present in the medium throughout the experiment. Therefore, it labels only structurally-intact membranes and when a membrane is disintegrated it will no longer label the malformed debris. Thus, these observations point to a scenario where major fractions of the distal membranes are completely detached from the main cell surface, gradually disintegrate, and lose their fluorescent labeling in the extracellular space between recently divided daughter cells.

To observe the same process at ultrastructural resolution we prepared dividing *S. turris* cells for TEM analysis. In short, synchronized cells were vitrified using high-pressure freezing and subsequently freeze-substituted and embedded in resin. This results in an optimal combination of near-to-native state preservation of subcellular structures paired with the ability of high-throughput TEM imaging at room temperature. The TEM images of *S. turris* cells prepared according to this procedure show exceptional preservation of the cellular environment and notably the SDV membranes and inorganic contents (Fig. S6).

We acquired hundreds of images from 227 cells at different stages of the cell cycle, and by categorizing and sorting these images, we reconstructed a timeline of valve formation and exocytosis in *S. turris* (Fig. 3). Cells with an SDV containing a growing valve were classified as being at the stage of valve formation (n=61, examples presented in Fig. 3 A-B’’). At these stages three lipid bilayers, the proximal and distal SDV membranes, as well as the plasma membrane, can be clearly distinguished. The distal SDV membrane is in very close proximity to the plasma membrane, with a distance of only 10-30 nm (Fig. 3 A’’ inset). During early valve formation, the SDV extends only as far as silica deposition has advanced (Fig. 3 A’’). As silica precipitation proceeds radially, the SDV expands with it. After formation of the porous base layer, the polygonal layer is formed on its distal side (Fig. 3 B-B’’, arrow). During the entire silicification process, the SDV membrane tightly delineates the growing valve, forming a highly confined space in which the silica morphology is under control of the SDV membrane. The valve reaches its final morphology while it is still fully enclosed in the SDV, we observed 23 cells with an intact SDV containing a fully mature valve (Fig. 3 C-C’’).

**Figure 3.**
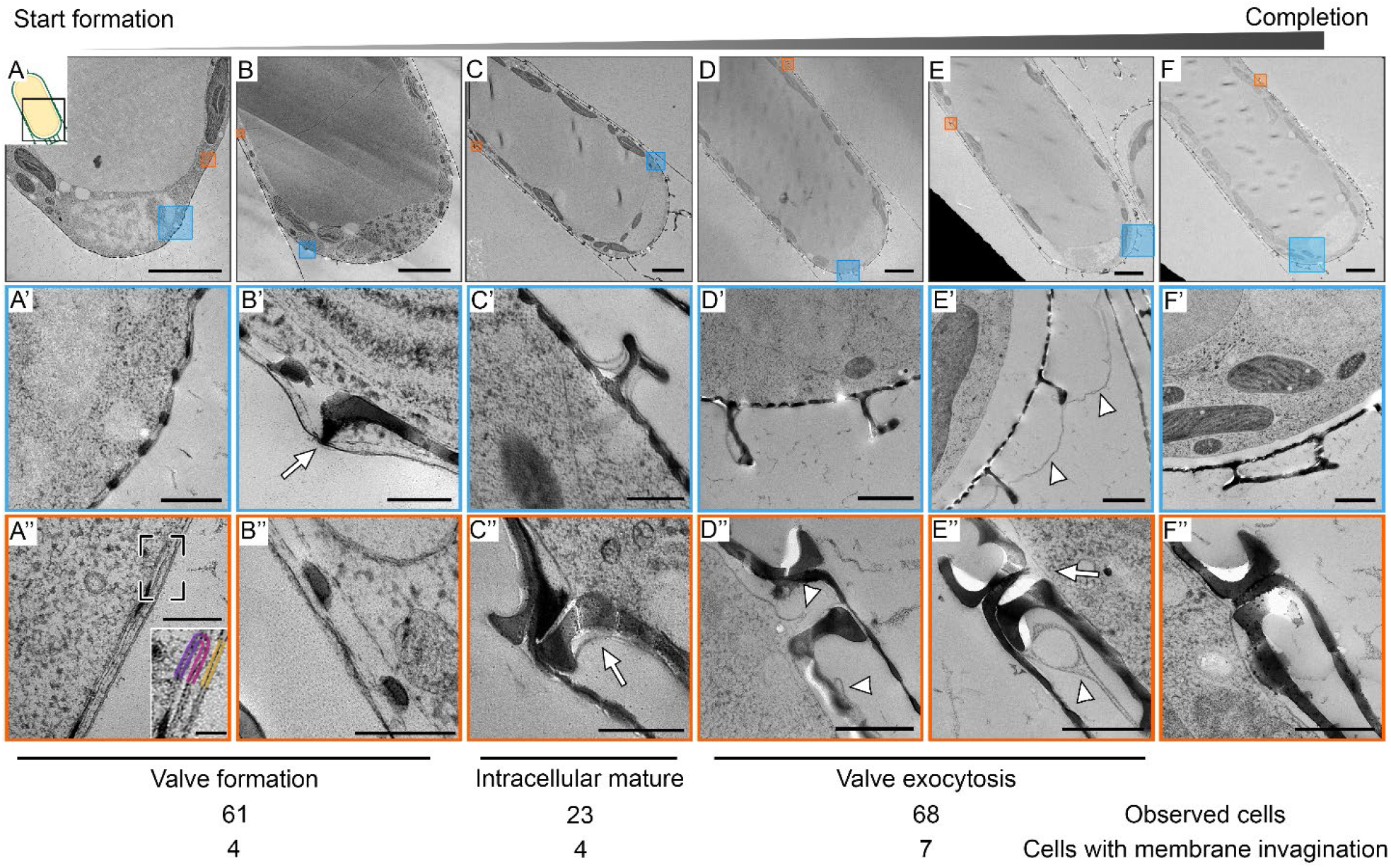
TEM images showing sequential stages of valve formation and exocytosis in *S. turris*. Top row shows a representative image of each stage, with higher magnification views of the boxed areas in the rows below. (**A**) Valve formation starts at the apex of the cell. (**A’**) Growing silica inside the SVD that is located directly under the plasma membrane. (**A’’**) The SDV ends at the growing silica edge. The inset shows the bilayers of the proximal (purple) and distal (pink) SDV and the plasma membrane (yellow) in higher magnification. (**B**) Later during valve formation, (**B’**) the polygonal layer (arrow) is formed on top of the base layer, and (**B’’**) the growing edge of the new valve reaches the rim of the parental valve. (**C**) Silicification of the new valve is completed while it is still fully enclosed in the SDV. Mature valves can be recognized by (**C’**) the fully formed and flattened polygonal layer and (**C’’**) pronounced hook shape at the valve rim (arrow). (**D**) At the onset of exocytosis, (**D’**) membranes still surround most parts of the new valve, (**D’’**) but they no longer form a complete enclosure (arrowheads). (**E**) Shortly after exocytosis, (**E’**) distal membranes lose their structural integrity (arrowheads), and (**E’’**) the plasma membrane is now continuous (arrow) under the new valve. (**F-F’’**) After completion of exocytosis, there are no traces left of the distal membrane remnants. Below the images, a table summarizes the number of cells observed in each developmental stage and the number of cases when membrane invaginations was present (see details in Fig. S7 and main text). Scale bars represent 5 μm (**A, B, C, D, E, F**), 1 μm (**D’, E’, F’**), 500 nm (**A’, C’, C’’, D’’, E’’, F’’**), 200 nm (**A’’, B’, B’’, C’** inset) and 50 nm (**A’’ inset**).

In images of 68 cells, we observed a mature valve that is surrounded by SDV membranes with some discontinuities, i.e. the SDV does not form a complete enclosure. Based on this structural information we define these cells as undergoing exocytosis. These membrane discontinuities can be very local, while the majority of the valve is still tightly enclosed by the SDV (Fig. 3 D-D’’). At a later stage, the membrane ultrastructure changes dramatically. The tight delineation of the distal SDV and plasma membrane around the valve loosens, and we observe deterioration of the structural integrity of the distal membranes (Fig 3. E-E’’). The membrane at the proximal side of the valve is now continuous with the plasma membrane under the parental valve (Fig. 3 E’’ arrow). At the end of the exocytosis process, the new valves can be recognized due to their position under the parental girdle bands, while no visible traces of the membranes are seen outside the cytoplasm (Fig. 3 F-F’’).

We analyzed the microscopy data of the membrane proximal to the newly formed valves in order to identify possible signs of endocytic membrane retrieval that might be associated with recycling the SDV membranes. We observed 7 occasions of membrane invaginations in cells at the exocytosis stage. The size of these invaginations varies from 0.5 μm to 3 μm (Fig. S7). However, such invaginations were detected at all stages of valve formation and exocytosis with similar occurrence (Χ^2^ _(df=2, N=152)_ = 2.23, p = .327), and thus cannot be correlated to membrane recycling after exocytosis. Overall, the TEM data is in agreement with the live-cell imaging, suggesting that during valve exocytosis the distal membranes disintegrate extracellularly without signs of retrieval, while the only visible membrane proximal to the new valve is the proximal SDV membrane.

In order to visualize the cellular organization as close as possible to the native state, we acquired cryo-ET data from *T. pseudonana* cells undergoing valve exocytosis. The smaller size of these cells makes them suitable for the needed sample preparation steps for imaging with cryo-ET. In short, we vitrified synchronized *T. pseudonana* cells using plunge freezing, prepared thin lamellae using a focused ion beam and then collected electron tomograms under cryogenic conditions (30, 34). Out of the 59 pairs of daughter cells that were imaged, 15 were during or shortly after valve exocytosis, i.e. before exocytosis of the first set of girdle bands. Four of those were during the early stages of exocytosis, with the new valve only partly covered by SDV membranes (Fig 4 A-D).

**Figure 4.**
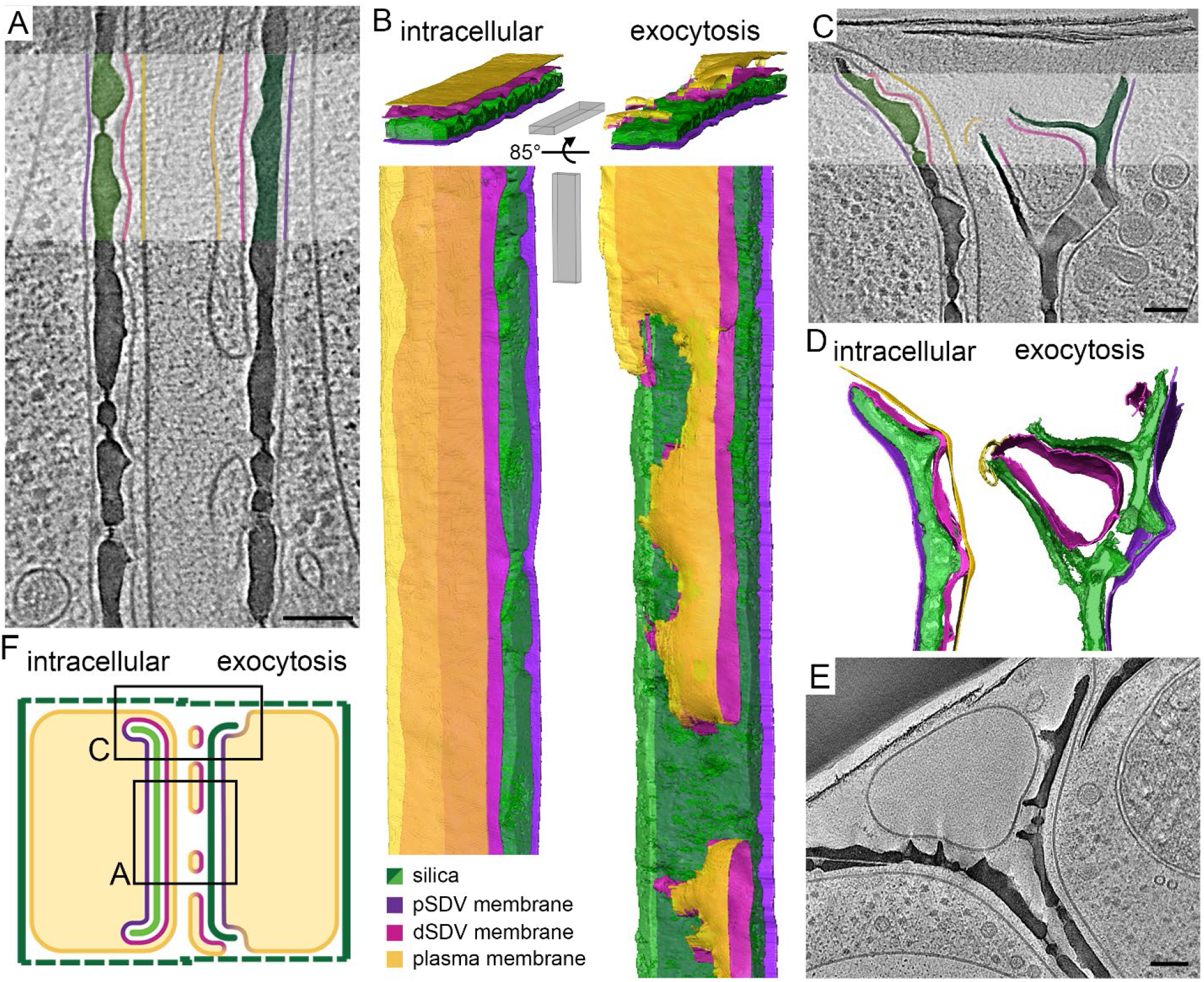
In-cell cryo-ET shows the native-state anatomy of valve exocytosis in *T. pseudonana*. (**A, C, E**) Slices through reconstructed 3D volumes; some features in the highlighted rectangles are artificially colored according to the color code in B. (**B, D**) Three-dimensional surface rendering of segmented volumes. (**F**) Schematic showing the proposed valve exocytosis mechanism of diatoms. **A** and **C** show different locations on the same daughter cells, indicated in **F**. Scale bars: 100 nm, slice thickness 1.15 to 5.7 nm. See also Movies S2-3.

Figure 4 A and C show tomograms of different locations of the same pair of slightly unsynchronized daughter cells, the left one shortly before exocytosis and the right one during exocytosis. The valve on the left side is still completely enclosed within the SDV, while the valve on the right side is already exposed to the extracellular space through discontinuities in the coverage by the distal membranes. In both cells, three lipid bilayers are discernible. In the daughter cell before exocytosis the expected arrangement of membranes is seen: the plasma membrane covers the whole cell and closely underneath the SDV membranes fully surround the valve (Fig. 4 A-B, left side). However, during exocytosis the distal membranes have fused at multiple sites, forming a network of flat membrane sacs, making the membrane on the proximal side of the valve the outermost boundary of the cell (Fig. 4 A-B, right side). A similar situation is observed at the valve periphery, where unconnected membranous structures are seen at the distal side of the fultoportula (Fig. 4 C-D, right side).

In eight pairs of cells that were at later stages of exocytosis we observed large membranous vesicles positioned between the recently exocytosed valves of two daughter cells and their girdle bands (Fig. 4 E). In the three remaining pairs of cells with recently exocytosed valves we did not observe any vesicles or membrane remnants between the two daughter cells. However, in those three cells the parental girdle bands were not enclosing the daughter cells, pointing to a later stage of the cell cycle, by which all extracellular debris had been degraded. Notably, none of the imaged cells contained budding endocytic vesicles or a sign for the formation of a new membrane underneath the valve. Therefore, the cryo-ET data suggest that *T. pseudonana* uses the same exocytosis mechanism that we inferred from *S. turris*, where the proximal SDV membrane is repurposed as a new plasma membrane and the distal membranes disintegrate extracellularly (Fig. 4 F).

## Discussion

Extrusion of the rigid cell walls of diatoms is an exceptional exocytosis event, difficult to reconcile with known exocytosis mechanisms. Our observations, of detached membrane patches in the extracellular space of two diatom species acquired with three microscopy techniques, are incompatible with classical exocytosis mechanisms. The structural characteristics of these unconnected and loose membranes after valve formation is in sharp contrast to the SDV ultrastructure during silica growth, which forms a highly confined space that is pivotal in controlling the shape of growing silica structures (35–37). Therefore, we suggest that exocytosis of mature valves occurs via a unique mechanism in which the old plasma membrane and distal SDV membrane are discarded in the extracellular space and the proximal SDV membrane takes the role of a new plasma membrane (Fig. 4F, Fig. S2). This scenario is in agreement with previous observations of rapid exchange of SDV proteins with the plasma membrane in live-cell studies of *T. pseudonana* (2, 38).

This scenario is also supported by the fact that none of the cells in our datasets show signs of processing or recycling of the SDV membranes, or the formation of a new plasma membrane under the valve. Nevertheless, we are aware that our live-cell imaging might not have the spatial resolution to detect such events, and that electron microscopy only gives information on random snapshots in time and space. For these reasons, we conducted a statistical analysis to determine the probability that exocytosis does involve the endocytic recycling of the SDV membrane. The tested scenario was that the ∼1μm invaginations observed in *S. turris* are actually part of such endocytic retrieval process (Fig. S7). If this is the case, it will require the recycling of ∼1300 such invaginations. We ran a statistical simulation, assuming a 10 seconds time window for an invagination to develop and be visible in an image, and a total 30 minutes for valve exocytosis. The simulation also addresses the probability to ‘catch’ this event in a random TEM thin slice (see supplementary material for simulation description). Running the simulation for 10,000 times showed that in more than 99.5% of the cases, the simulated scenario included the detection of more than 7 such events when 68 observations were made (corresponding to the 68 actual observations, Fig. 3). Therefore, we can rule out the option of internalization of these membranes through compensatory endocytosis with a probability of p<0.001, according to the permutation test.

The proposed scenario suggests that upon exocytosis the proximal SDV membrane becomes the new plasma membrane. As the SDV and plasma membranes carry out specialized, different tasks, it is likely that both require unique lipid and protein compositions, which would need to be adjusted after fusion (39). The finding that the distal membranes disintegrate extracellularly is surprising, as it seems wasteful to discard such a large amount of membranes each cell cycle. The low affinity of FM4-64 to degrading debris precludes the ability to follow the fate of these membranes (32). Nevertheless, it is important to note that the girdle bands form a quasi-enclosed compartment around the newly formed valves, possibly slowing down diffusion away from the cell and facilitating the uptake of the membrane debris for cellular use after disintegration. This can conceivably minimize the energetic costs of such process.

The exocytosis of large content was also investigated in other organisms, demonstrating mechanisms that differ from the classical secretory pathway of numerous small vesicles that fuse with the membrane (40). One alternative is the exocytosis of giant vesicles in the *Drosophila* salivary glands that squeeze out their viscous cargo through a pore by crumpling of the vesicular membrane, followed by membrane recycling (41). Another alternative, the acrosome reaction, surprisingly shares similarities with our proposal for diatom valve exocytosis. In this process, the plasma membrane and distal acrosomal membrane fuse at several locations forming vesicles that are dispersed in the environment while the proximal acrosomal membrane remains intact and becomes the new boundary that separates the spermatozoa from the environment (42, 43).

To conclude, this work proposes a unique mechanism for the exocytosis of diatom silica, which involves two remarkable events. First, the repurposing of an organelle membrane into a plasma membrane and second, large-scale disintegration of membranes. This mechanism is shared by two model diatom species, indicating that this might be a general mechanism. With the toolbox for genetic research in diatoms growing, it will soon be possible to investigate the protein machinery that is involved in the regulation of this event, and its relation to classical exocytosis.

## Acknowledgements

We thank Anne Jantschke, Nathalie Pytlik, and Eike Brunner from TU Dresden for providing cultures. We thank Simon Michaeli from the Volcani Institute (ARO) for his contribution in finding the right live-cell imaging platform. Live-cell imaging was conducted at the de Picciotto Cancer Cell Observatory In Memory of Wolfgang and Ruth Lesser. This project has received funding from the European Research Council (ERC) under the European Union’s Horizon 2020 research and innovation programme (grant agreement No. 848339). D. dH. was supported by the Sustainability and Energy Research Initiative (SAERI) of Weizmann Institute of Science.

## Data availability

All source data generated or analyzed during this study will been deposited to Dryad.

## Supplementary information

### Materials and methods

#### Cell cultures

*S. turris* was isolated from the North Sea in 2004 and provided by the group of Prof. Eike Brunner, TU Dresden. Diatom cultures were maintained in natural Mediterranean seawater that was filtered, its salinity corrected to 3.5% and supplemented with f/2 nutrient recipe, and for *Thalassiosira pseudonana* (CCMP1335) supplemented with 330 μM silicic acid. Cultures were maintained at 18°C under 16/8 hours light/dark cycles.

#### Cell cycle synchronization

For synchronization of the cell cycle, 5 ml of a mature *S. turris* cell culture was used to start a new 50 ml culture. After 48 hours of growth under the normal 16L/8D cycle, the culture was placed in darkness for 20 hours. After 20 hours of darkness, the culture was again exposed to light. *T. pseudonana* synchronization was done using Si starvation, as previously described (7). To maintain the culture in an exponential growth phase they were grown under 12/12 hours light\dark cycles and diluted (1:10) into fresh medium every other day during the week before culture synchronization. To induce Si starvation, 100 ml aliquots of culture were centrifuged at 3000 g for 10 min and re-suspended in Si-free artificial seawater or filtered seawater; this step was repeated three times. The cultures (∼0.5 million cells/ml), were then maintained in dark for 12 hours under agitation in a Si free medium to arrest the cell cycle. Then cultures were transferred to continuous light for an additional 4 hours of Si starvation. At the end of the Si starvation period, cells were concentrated to about 10 million cells/ml and Si was replenished to 330 μM. To track the formation of new silica, PDMPO [2-(4-pyridyl)-5-((4-(2-dimethylaminoethyl-amino-carbamoyl)methoxy)-phenyl) oxazole] (ThermoFisher Scientific, USA) was added. PDMPO fluorescence was monitored by imaging the cultures with an epifluorescence microscope (Nikon Eclipse Ni-U, ex: 365 em: 525). Highest amount of dividing *S. turris* and *T. pseudonana* cell were counted after 9 and 3 hours, respectively.

#### Sample preparation for SEM

Cells were prepared for SEM using critical point drying (CPD). Cells were fixed in a solution of 2% Glutaraldehyde and 4% Paraformaldehyde in artificial seawater for 1 hour at room temperature while shaking. After 3 washes with deionized water (Milli-Q® IQ 7003 Ultrapure Lab Water System, Merck), the cells were dehydrated by washing in a graded series of ethanol. The final wash was done in 100% anhydrous ethanol overnight. The dehydrated samples were then dried in a critical point dryer using liquid CO_2_ as transitional fluid. Dried cells were placed onto a conductive carbon tape on an aluminium stub.

#### SEM imaging

Samples were sputter-coated with 2.5 nm (*T. pseudonana*) or 4 nm (*S. turris*) iridium (Safematic) and imaged with an Ultra 55 FEG scanning electron microscope (Zeiss, Germany), using 3-5 kV, aperture size 20-30 um and a working distance of about 3 mm.

#### Live-cell imaging

For single-cell time-lapse imaging, a culture was synchronized and stained with PDMPO (330 μM). About 8 hours after the end of light starvation, when most cells had gone through cytokinesis, an aliquot of 100 μl was taken and FM4-64 (ThermoFisher Scientific, USA) was added to a final concentration of 4-8 μM to stain the membranes. After adding the membrane dye, a 20 μl drop was mounted on a microscope slide and covered with a glass coverslip, using dental wax as spacer. The samples were visualized using a Leica TCS SP8-STED confocal microscope equipped with a HCS PL APO 86x/1.20W motCORR objective. FM4-64 and chlorophyll autofluorescence were acquired by white-light laser using 550 nm laser line (6% laser power) and 650 nm laser line (7% laser power), respectively. PDMPO fluorescence was acquired using a 405 nm laser (5% laser power). HyD-SMD detectors were used for PDMPO and FM4-64 with emission collection width set to 474-530 and 608-640 nm, respectively. Chlorophyll autofluorescence emission was collected using a HyD detector with 741-779 nm detection width and the transmission channel was detected with a PMT detector. The cells were imaged for 2 to 4 hours at intervals of 3 to 60 minutes. Images were analysed using Leica Application Suite X. In total, time lapses from over 50 cells were collected over the entire period.

#### Sample preparation for TEM

*S. turris* cells were cryo-fixed, using high pressure freezing (HPF), followed by freeze substitution (FS), according to previously published protocols (29, 44). Synchronized cells were collected on a 5 μm filter membrane and transferred to an Eppendorf tube using 200 μl of seawater. After letting the cells sediment for 10 minutes, 2 μl aliquots were pipetted into aluminium discs (Wohlwend GmbH, Sennwald, Switzerland) and directly loaded into a Leica ICE high pressure freezing machine (Leica Microsystems GmbH, Wetzlar, Germany). Concentrated diatom samples were vitrified in liquid nitrogen (−192 °C) at 210 MPa (2048 bar). Vitrified samples were stored in liquid nitrogen until freeze-substitution in an EM ASF2 (Leica Microsystems GmbH, Wetzlar, Germany). Vitrified water was substituted with an organic solvent, 100% anhydrous acetone, at (−90 °C). Then, acetone was supplemented with chemical fixatives (0.2% uranyl acetate and 0.2% osmium tetroxide) to enhance cross-linking and contrast of cellular structures. The samples were immersed in the solution for 48 hours at −90 °C and then allowed to gradually warm for 24 hours to -20°C, and then in one hour to 0°C. After three washes in acetone, the acetone was replaced with Epon (Agar Scientific Ltd, Stansted, U.K.) using gradient concentration mixtures (10%, 20%, 30%, 40%, 60%, 80%, 100% Epon in acetone), twice a day at room temperature. The sample in 100% Epon was hardened at 70 °C for 72 h. Ultrathin sections of 70 nm were sliced using an ultra-microtome (Ultracut UCT, Leica Microsystems GmbH, Wetzlar, Germany) equipped with a diamond knife (Ultra 45°, Diatome Ltd, Nidau, Switzerland). The sections were picked up onto copper TEM grids, coated with carbon film. The sections were then post-stained by placing them onto a drop of lead citrate solution. After three minutes of staining, the grids were washed three times in drops of water and then dried by blotting onto whatman filter paper. In total, we prepared 16 frozen samples that were further processed using freeze substitution during 3 different cycles, and subsequently imaged hundreds of cells.

#### TEM imaging

The TEM samples were imaged with a Tecnai Spirit TEM (FEI, Eindhoven, Netherlands) operated at 120 kV and equipped with Gatan Oneview 4 k × 4 k camera (Gatan Inc., Pleasanton, U.S.A.).

#### Plunge freezing

Synchronized *T. pseudonana* cells were vitrified by plunge-freezing on glow discharged 200 mesh copper R2/1 holey carbon film grids (Quantifoil Micro Tools GmbH, Grossloebichau, Germany). In a Leica EM GP (Leica Microsystems GmbH, Wetzlar, Germany), 1 μl of artificial seawater was pipetted on the copper side in order to enhance media flow to the blotting paper and 4 μl of cell suspension at 7-13×10^6^ cells/ml was pipetted on the carbon side. The grids were blotted for 6 seconds from the back side of the grid before they were plunged into a liquid ethane bath cooled by liquid nitrogen.

#### Cryo-FIB milling

Vitrified cells were milled to thin lamellae with the Zeiss Crossbeam 550 FIB/SEM dual beam microscope (Zeiss, Germany). The grids were coated with organometallic platinum by an *in situ* Gas Injection System. The lamellae were milled at a 12º tilt relative to the grid plane with the rough milling (to 1 μm thickness) involving 2 steps using the Ga^+^ beam at a current of 750 pA and 300 pA. After rough milling all lamellae, they were thinned to 200 nm at a current of 50 pA.

#### Cryo-electron tomography and volume rendering

Cryo-electron tomography data were collected from 59 pairs of cells. The tilt series were acquired using a Titan Krios G3i TEM (Thermo-Fisher Scientific, Eindhoven, The Netherlands), operating at 300 kV. Tilt series were recorded on a K3 direct detector (Gatan Inc., Pleasanton, U.S.A) installed behind a BioQuantum energy filter (Gatan Inc., Pleasanton, U.S.A), using a slit of 20 eV. All tilt series were recorded in counting mode at a nominal magnification of 33,000x, corresponding to a physical pixel size of 0.26 nm, using the dose-symmetric scheme starting from the lamella pre-tilt of -12º and with 2º increments (45). Tilt series appearing in Fig. 4 A and C were taken at 3 μm defocus and using a Volta Phase Plate inserted. The tilt series range was between -60° to 50°. Tilt series appearing in Fig. 4 E was taken at 7 μm defocus, an objective aperture of 100 μm inserted, and -66° to 48° tilt range. Tilt series were acquired using an automated low dose procedure implemented in SerialEM with a total dose set to ∼100e-/Å2 (46). The tomograms were reconstructed using IMOD software v. 4.9.12 (47). Amira software v2021.2 was used for segmentation (Thermo-Fisher Scientific, Eindhoven, The Netherlands). Membranes were segmented using the membrane enhancement filter module and manually refinement (48).

#### Statistical simulations and analyses

To test whether our data indeed indicate a novel exocytosis strategy rather than classical exocytosis we ran simulations and 10000 permutations. In order to do so, we estimated how many vesicles would have to be endocytosed to re-internalize a membrane equivalent in size to the mature SDV of *S. turris* and simulated how many such vesicles should have appeared in our dataset.

**SDV membrane surface area** (excluding the membranes around linking processes) ≈ Total plasma membrane surface area ≈ surface area of a capsule

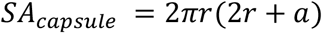

*S. turris* cells are 20 to 25 μm in diameter and 60 μm in length, so the surface area for an average sized *S. turris* cell:

r =11 μm

a =40 μm

SA_capsule_ ≈ 4000 μm^2^

### Membrane surface area of a vesicle

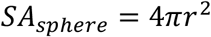

In our data we observed membrane invaginations of 0.5 to 1 μm. Membrane surface area of the invaginations seen in *S. turris*:

r = 0.25 to 0.5 μm

SA_sphere_ = 0.78 μm^2^ to 3.14 μm^2^

### Number of vesicles to be endocytosed

If all of the membrane is endocytosed by vesicles of 0.5 μm:

4000/0.78 = 5128 vesicles

If all of the membrane is endocytosed by vesicles of 1 μm:

4000/3.14 = 1274 vesicles

### Chance of catching an endocytic vesicle in a TEM section

The chance of detecting a spherical vesicle of diameter x in a random slice going through the round cross section of a cell with diameter y is

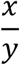

The chance to detect a vesicle of 1 μm in a TEM section of an *S. turris* cell of 25 μm is

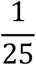

The chance to detect a vesicle of 0.5 μm in a TEM section of an *S. turris* cell of 25 μm

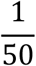

### Duration of an endocytosis event for a 0.5-1 um vesicle

Studies of time resolved endocytosis indicate that a single endocytosis event of a vesicle with a diameter of 0.5 μm should take at least 10 seconds. (49–52)

### Total duration of the exocytosis/endocytosis process

Based on our data, we surmise that the recycling process should be finished in 30 minutes.

Using the parameters described above we find that if compensatory endocytosis would take place, we should see more than 7 vesicles in >99.5% of the cases.

The simulation was run in R, v. 4.1.2.

A histogram showing the frequency of simulations that detected each number of endocytic vesicles during 68 observations. The two histograms show the two different vesicle sizes that were simulated.

**Figure S1.**
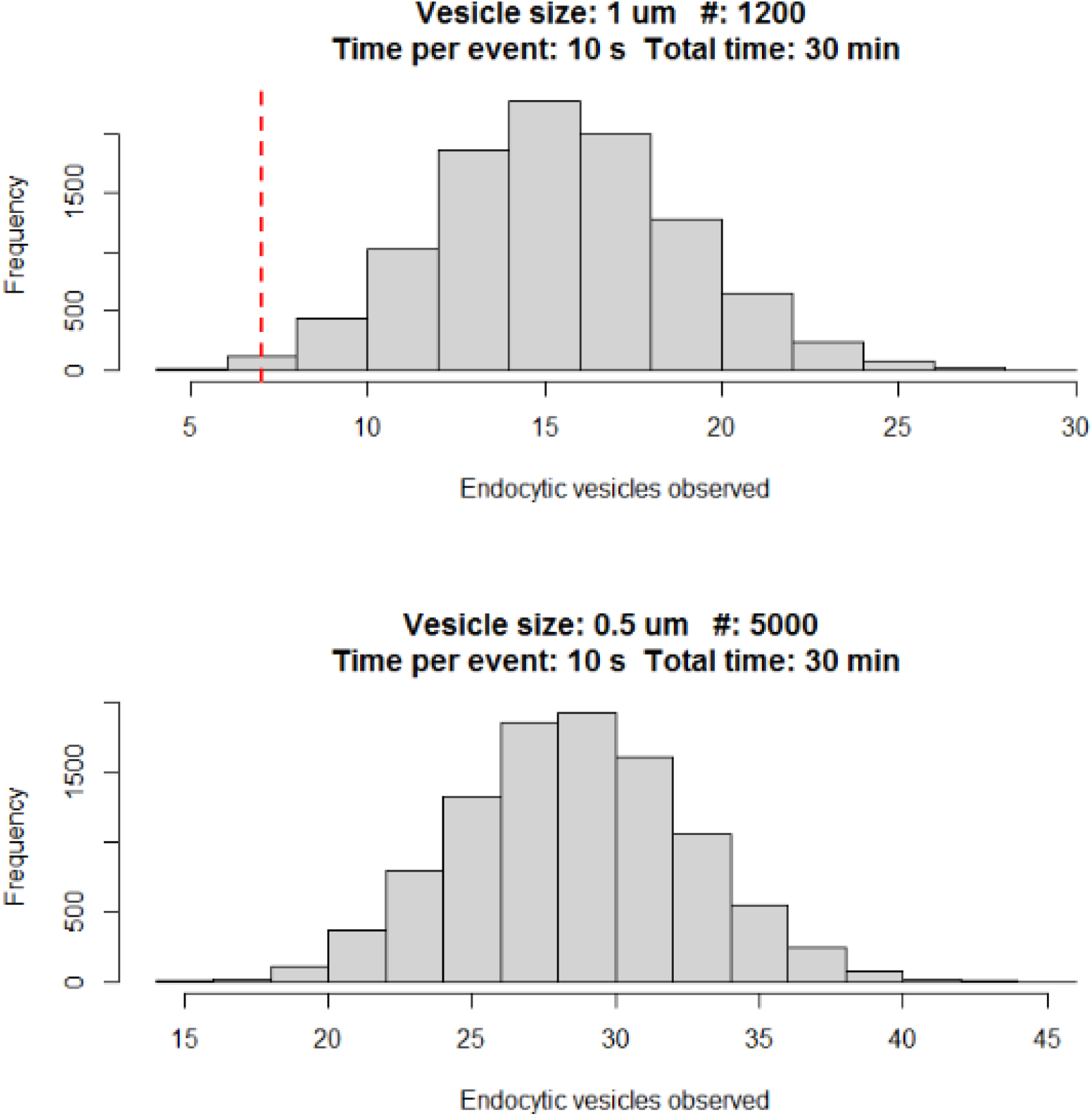
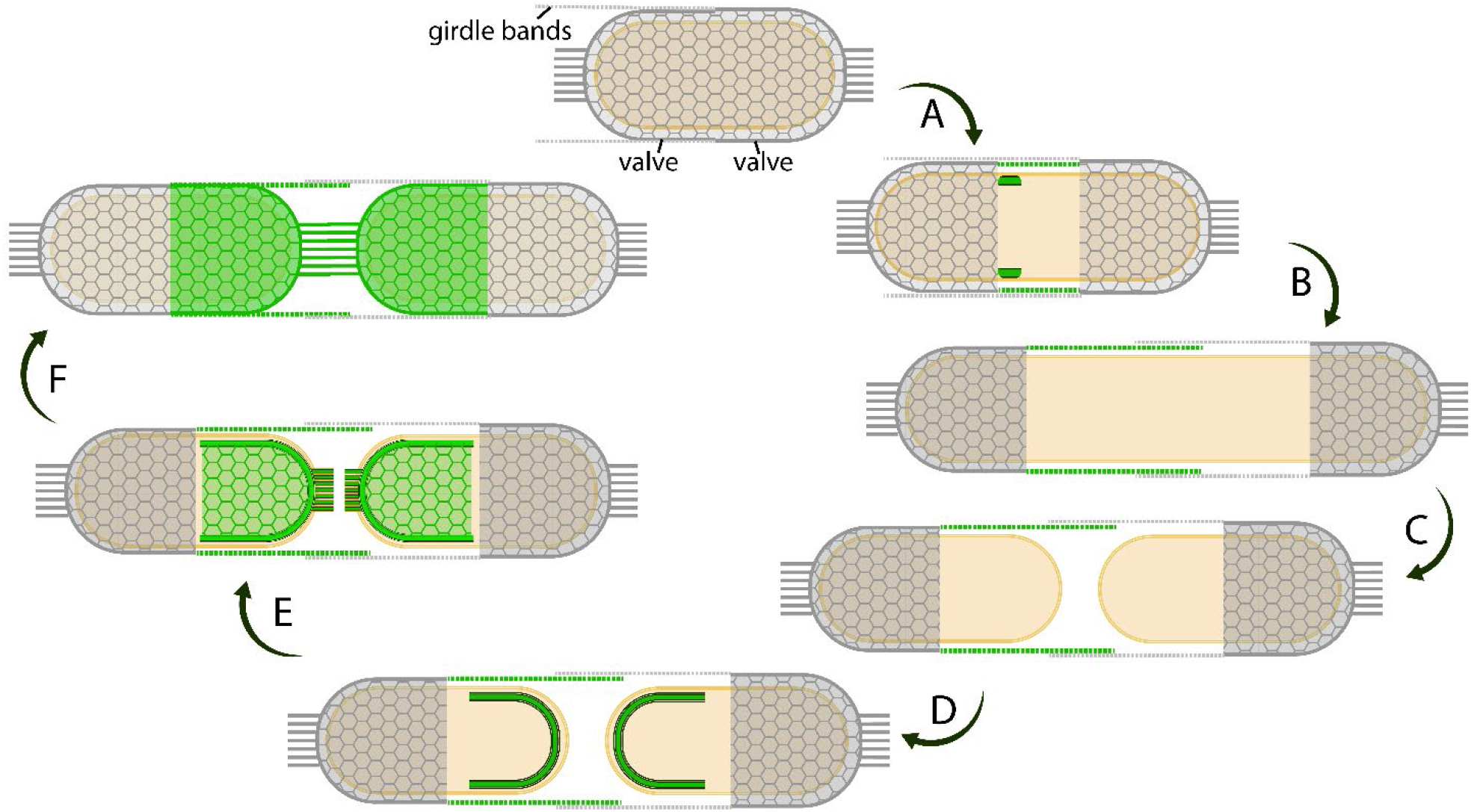
Cell cycle of *S. turris*. New silica elements, formed during the current cell cycle are colored green. Dashed lines represent girdle bands. (**A**) Start of a cell cycle with a protoplast enclosed by the two parental valves, the valve rims are directly adjacent to each other. The older valve is on the right, and the younger valve on the left. The younger valve is covered under the girdle bands of the older valve. (**B**) Protoplast starts growing in preparation for the next cell division. As the silica valves are rigid and not elastic, growth is only possible in longitudinal direction by pushing apart the two valves. New girdle bands (green) are formed in individual SDVs and added to the rim of the younger valve. (**C**) After the protoplast has reached its final length, the cell undergoes cytokinesis, resulting in two daughter cells enclosed in the parental cell wall. (**D**) Each daughter cell inherits one half of the parental cell wall and starts forming a new valve, inside an SDV, shortly after cell division. (**E**) The new valves are completed with attachment of the linking extensions, after which they are exocytosed. (**F**) Both daughter cells are now ready to continue to the next cell cycle.

**Figure S2.**
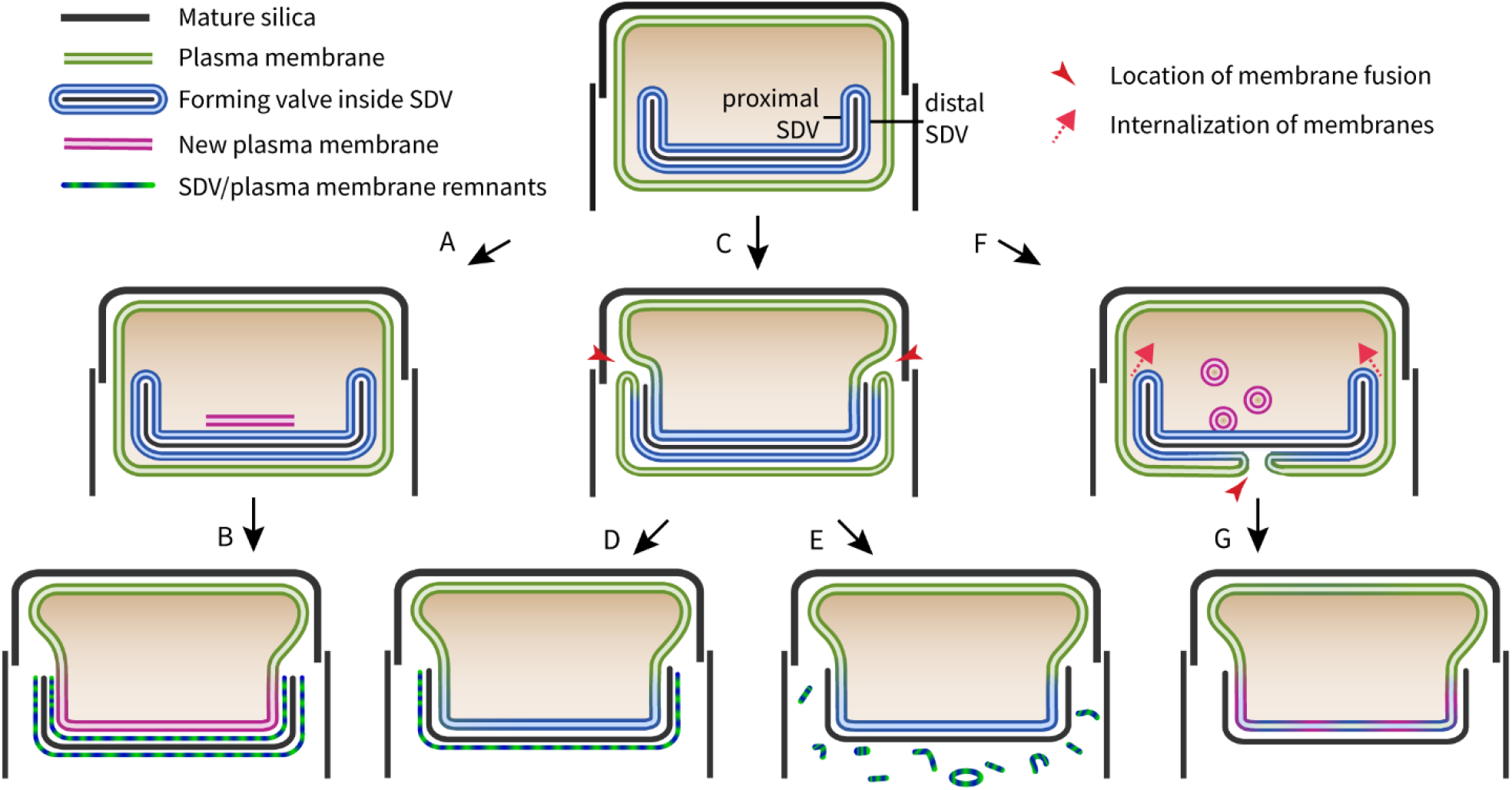
Hypotheses for valve exocytosis in diatoms. Early ultrastructural studies brought forward few models for silica cell wall exocytosis in diatoms that are reproduced here (14, 16– 23). One scenario suggests that the newly formed valve is externalized through formation of a new plasma membrane underneath it (**A**). The SDV membrane and the original plasma membrane around the new valve remain in place, forming an organic layer around the silica (**B**). An alternative is that the valve is externalized as a result of localized fusion of the SDV and plasma membrane at the valve edge, such that the proximal part of the SDV membrane becomes the new plasma membrane (**C**). The distal part of the SDV membrane and the original plasma membrane were suggested to either shed off to the environment (**E**) or remain as an extracellular layer protecting the silica (**D**). The last model suggests localized membrane fusion at the apical part of the valve, followed by pulling of the distal SDV and original plasma membrane back into the cytoplasm at the valve edge (**F**). In this last model, which is in accordance with classical exocytosis, the composition of the proximal SDV membrane is altered for its new function as the plasma membrane (**G**). Our study support model C+E as the scenario for valve exocytosis.

**Figure S3.**
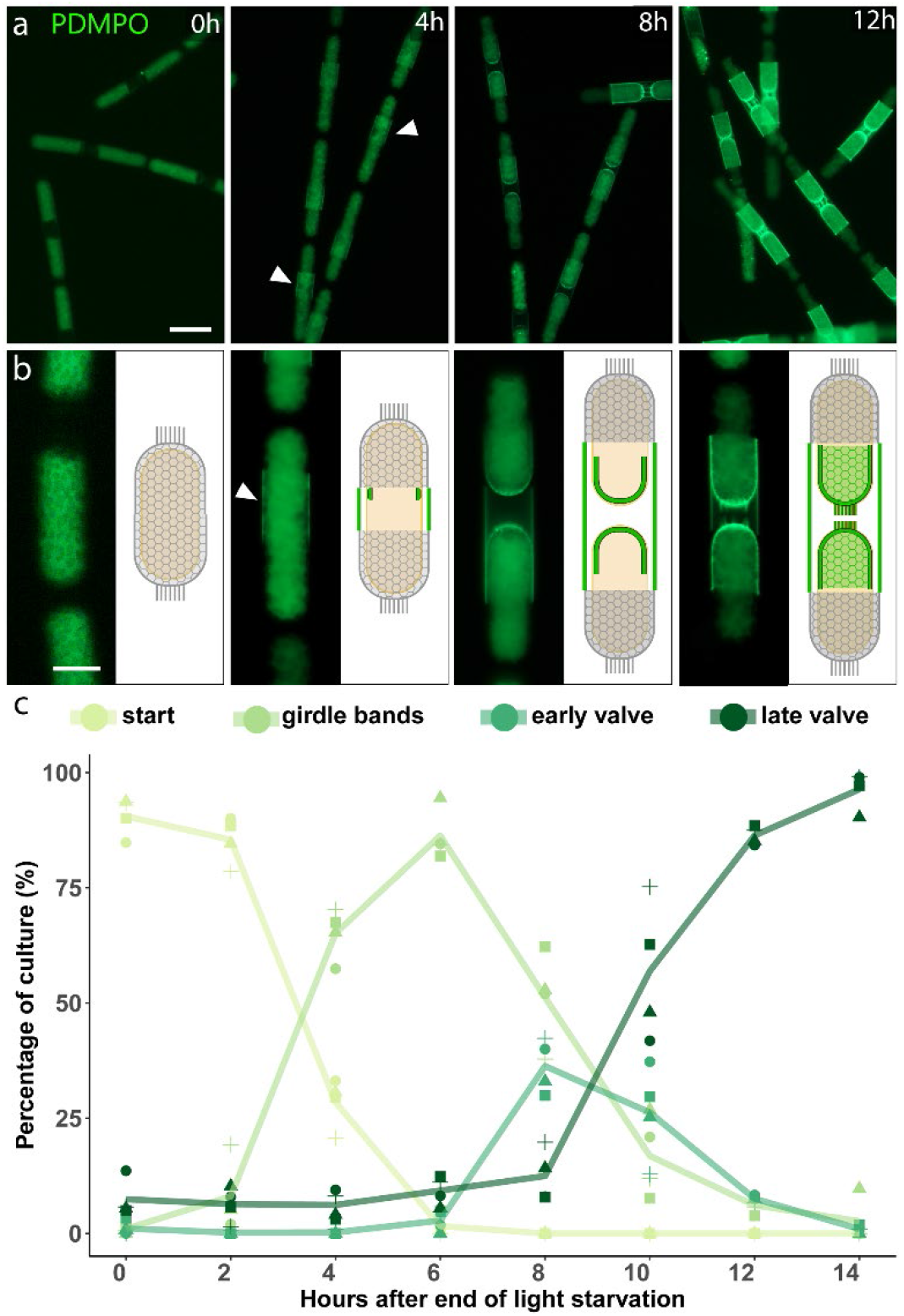
Light-induced synchronization of *S. turris* cell cycle. (**A**) Representative PDMPO fluorescence images of synchronized cultures. Elapsed time since the end of light starvation and addition of PDMPO is indicated in the image. Shortly after the end of light starvation, none of the cells have fluorescently labelled silica, only PDMPO fluorescence from within the vacuoles is visible. Four hours later, most cells have started forming girdle bands, visible as a fluorescent ring (arrowheads) around the cell. After eight hours, the majority of the culture is forming new, fluorescent valves. After twelve hours, nearly all cells have completed the formation of a new valve. Scale bar: 50 μm. (**B**) Key stages of cell wall formation: ‘start’, cells without PDMPO-labelled silica; ‘girdle bands’, cells during cell elongation and girdle band formation; ‘early valve’, after cell division, valve formation has started but is prior to growth of linking extensions; ‘late valve’, cells at later stages of valve formation when polygons and extensions are visible. Scale bar: 20 μm (**C**) Percentage of cells at key stages during synchronized growth. The lines show the mean of four independent experiments, indicated with symbols. The highest percentage of cells that are forming valves is reached around nine hours.

**Figure S4.**
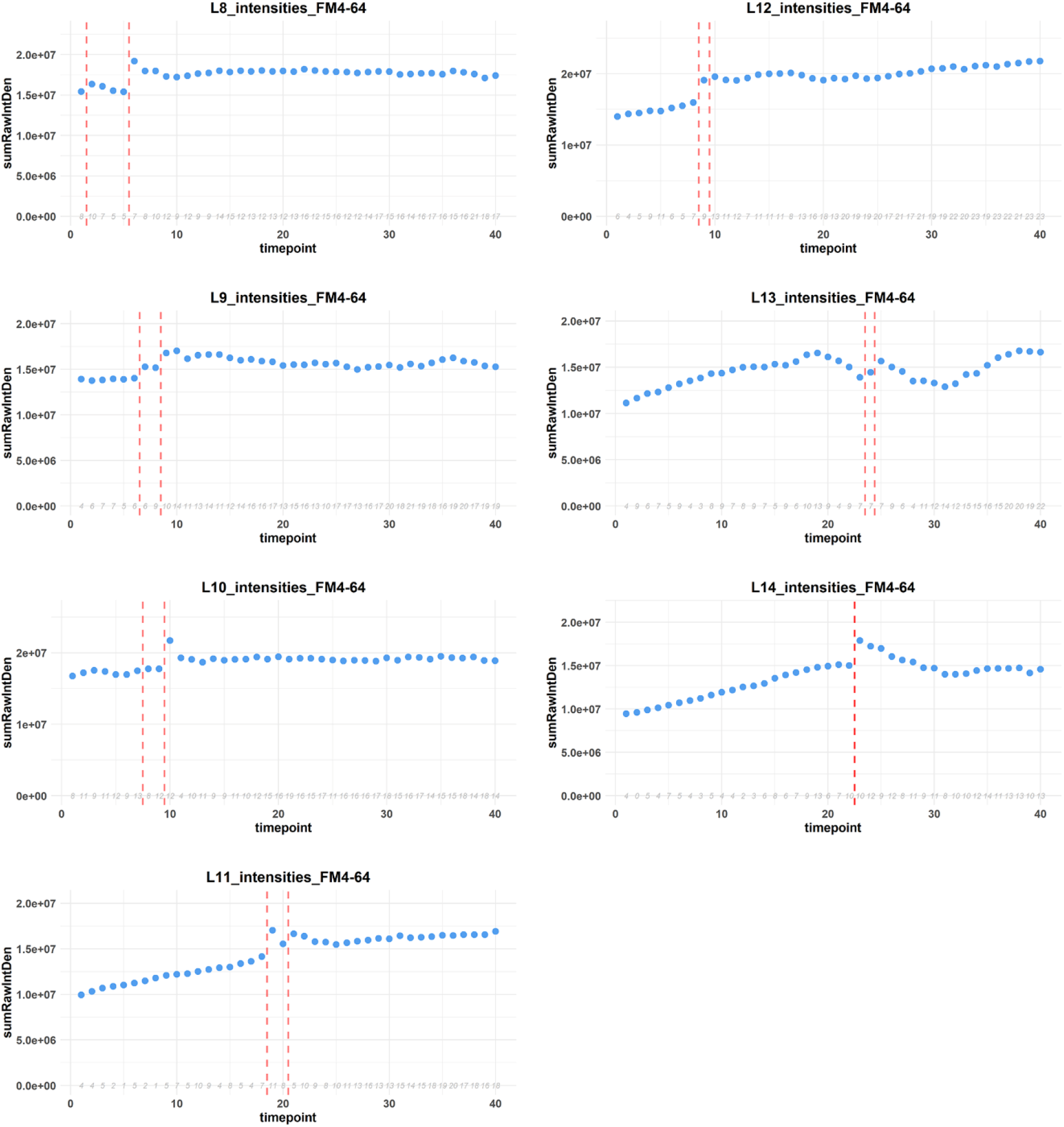
FM4-64 fluorescence intensity during live-cell imaging. Results from seven cells undergoing valve formation and exocytosis are shown. Total FM4-64 fluorescence intensity throughout time-lapses were measured using imageJ’s ‘raw integrated density (RawIntDen)’. This measure gives the sum of all pixel intensities in an image. Here we plotted for each time point the sum of RawIntDen of all slices in the z-stack. The red lines indicate where exocytosis happens, based on loosening of the hexagonal pattern. Thus, in 6 out the 7 cells (except for L13) a clear fluorescence increase is associated with the disappearance of the polygonal pattern. In L14 only one of the two daughter cells remained in the field of view. Grey italic numbers indicate for each time point the number of slices that contain one or more oversaturated pixels, the Z-stacks contain 40 to 44 slices overall.

**Figure S5.**
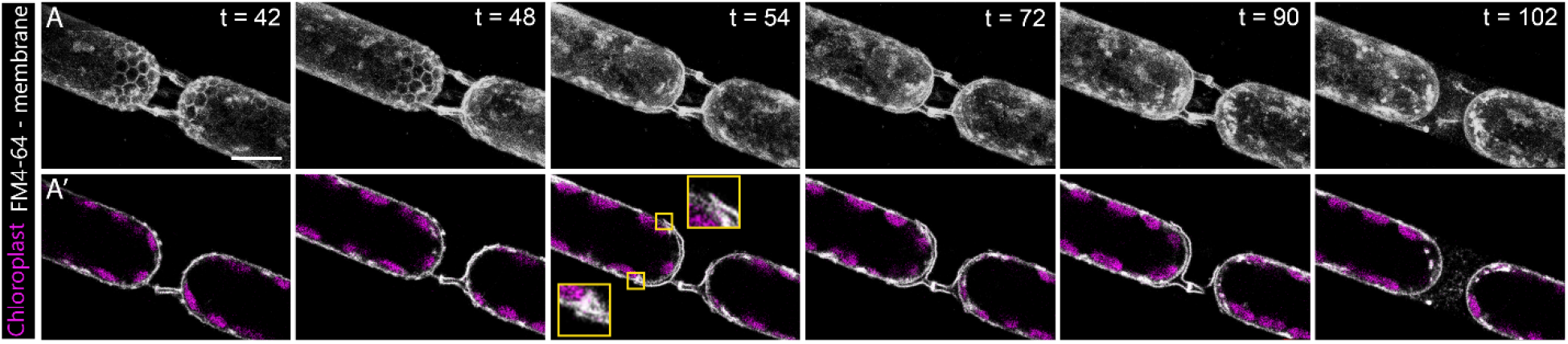
Distal membranes that undergo degradation are disconnected from the cellular membrane. Representative (**A**) maximum intensity projections and (**A’**) single z-slices of time-lapse confocal fluorescence images. Images show membrane staining (FM4-64) in white and chloroplasts autofluorescence in magenta. The elapsed time since the start of imaging (in minutes) is shown on the upper right corner. Exocytosis of the valve on the bottom right starts between t=42 and t=48. The valve on the top left is exocytosed between t=48 and t=54. In t=54 to t=90, distal membranes around the valve are not connected to the cellular membrane. At t=102 the remnants have largely disintegrated. Scale bar: 10 μm.

**Figure S6.**
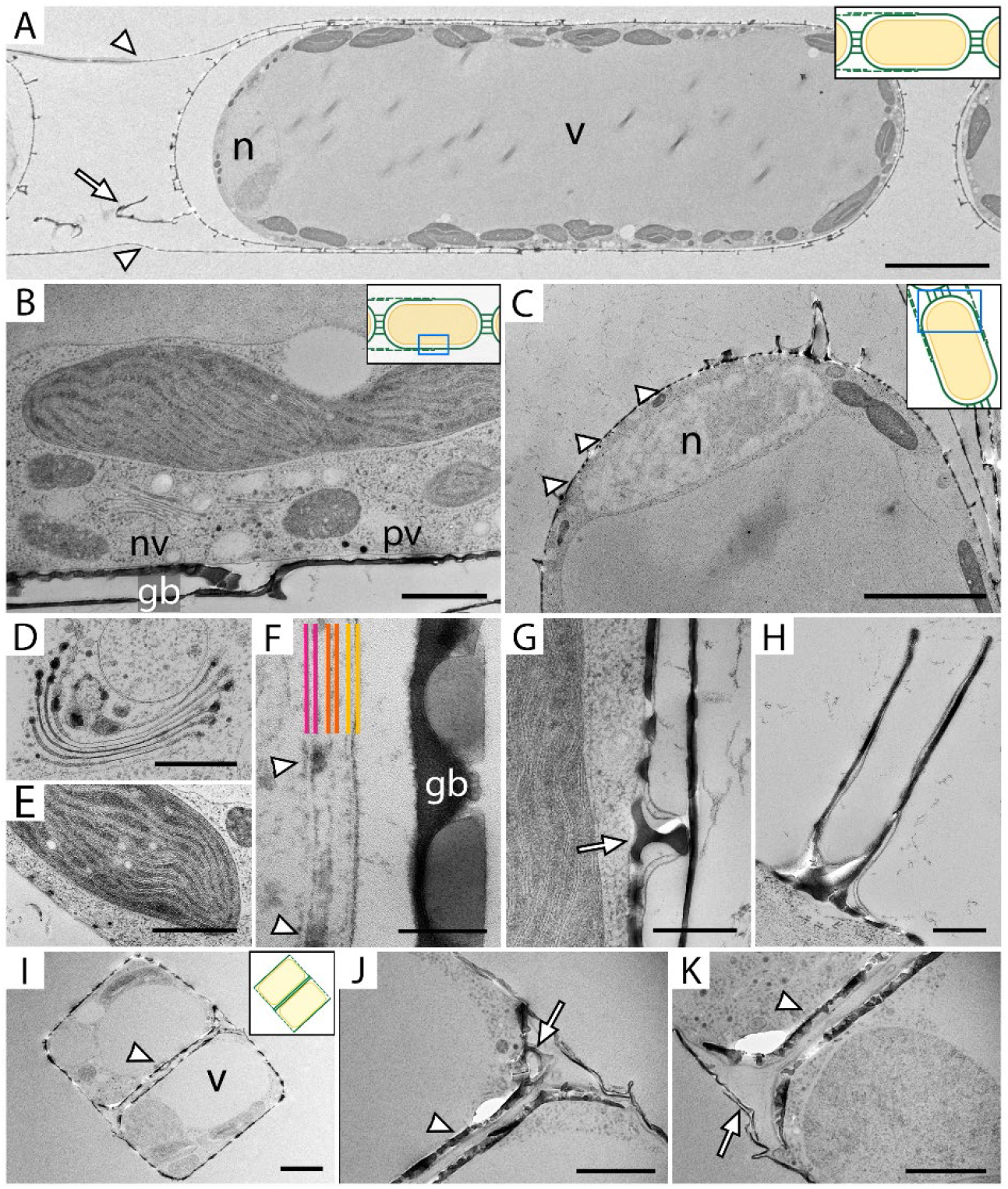
Ultrastructural details of diatoms revealed by TEM imaging. Insets indicate which part of the cell is displayed. (**A**) A cross section through an entire cell, the nucleus (n), and the large central vacuole (v) that is surrounded by darker chloroplasts are dominant. The silica cell wall (dark contrast) is clearly visible at the left side, where the protoplast is not directly adjacent to the cell wall due to slight plasmolysis. Arrowheads indicate girdle bands, and an arrow indicates a linking extension. Scale bar: 10 μm. (**B**) The meeting point of a new valve (nv), and parental valve (pv) with girdle bands (gb) attached to the latter. Scale bar: 1 μm. (**C**) Apex of a cell with the nucleus (n) located directly under the valve. Silica protrusions indicated with arrowheads are cross sections of the polygonal layer. Scale bar: 5 μm. (**D-H**) High magnification images of: (**D**) Golgi-body. Scale bar: 500 nm. (**E**) Chloroplast. Scale bar: 1 μm. (**F**) SDV with growing silica (arrowheads). Bilayers of the proximal SDV membrane (pink), distal SDV membrane (orange), and plasma membrane (yellow) can be distinguished. Scale bar: 100 nm. (**G**) Fully formed polygon (arrow) still inside an SDV. Scale bar: 500 nm. (**H**) Linking extension during its formation in an SDV. Scale bar: 500 nm. (**I**) Cross section of two *T. pseudonana* daughter cells with intracellular valves (arrowheads). Scale bar: 1 μm. (**J-K**) Higher magnification of the cells in **I**, with arrow in **J** indicating a fultoportula in the forming valve and arrow in **K** indicating parental girdle bands. Scale bars: 500 nm.

**Figure S7.**
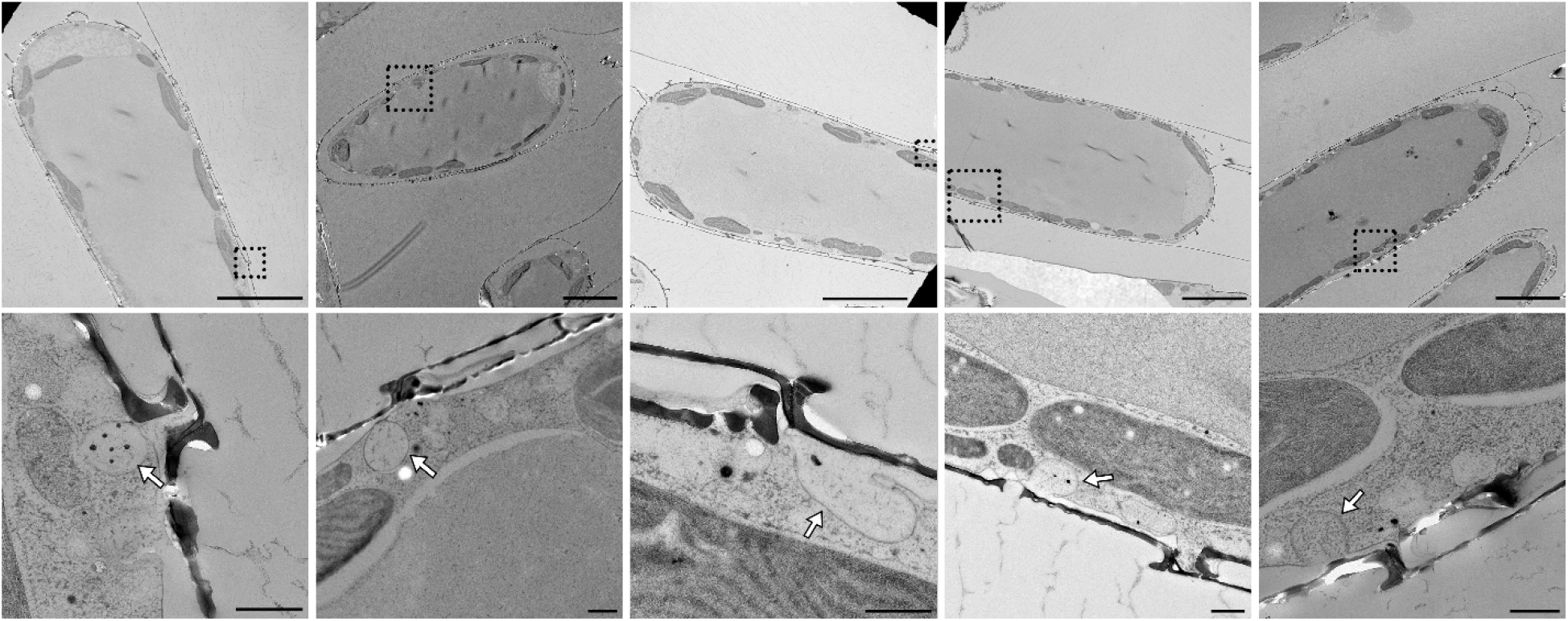
Instances of membrane invagination during valve exocytosis. Examples of cells that are undergoing valve exocytosis, i.e. have a mature valve that is exposed to the exterior but still surrounded by membrane (remnants). The bottom row shows magnified views of the boxed areas in the top row, arrows indicate membrane invaginations. Scale bars: 1 μm (top row) and 500 nm (bottom row).

**Movie S1**. Time-lapse imaging video of *S. turris* cell, stained with FM4-64, during valve formation and exocytosis. (MP4 2186 kb)

**Movie S2**. Animation of cryo-ET dataset of *T. pseudonana* shown in Figure 4 A. (MP4 51 Mb)

**Movie S3**. Animation of cryo-ET dataset of *T. pseudonana* shown in Figure 4 C (MP4 51 Mb)

